# Dysregulation of EMT Drives the Progression to Clinically Aggressive Sarcomatoid Bladder Cancer

**DOI:** 10.1101/388264

**Authors:** Charles C. Guo, Tadeusz Majewski, Li Zhang, Hui Yao, Jolanta Bondaruk, Yan Wang, Shizhen Zhang, Ziqiao Wang, June Goo Lee, Sangkyou Lee, David Cogdell, Miao Zhang, Peng Wei, H. Barton Grossman, Ashish Kamat, Jonathan James Duplisea, James Edward Ferguson, He Huang, Vipulkumar Dadhania, Colin Dinney, John N. Weinstein, Keith Baggerly, David McConkey, Bogdan Czerniak

## Abstract

The sarcomatoid variant of urothelial bladder cancer (SARC) displays a high propensity for distant metastasis and is associated with short survival. We report a comprehensive genomic analysis of 28 cases of SARCs and 84 cases of conventional urothelial carcinomas (UCs), with the TCGA cohort of 408 muscle-invasive bladder cancers serving as the reference. SARCs showed a distinct mutational landscape with enrichment of *TP53, RB1, and PIK3CA* mutations. They were related to the basal molecular subtype of conventional UCs and could be divided into epithelial/basal and more clinically aggressive mesenchymal subsets based on TP63 and its target genes expression levels. Other analyses revealed that SARCs are driven by downregulation of homotypic adherence genes and dysregulation of cell cycle and EMT networks, and nearly half exhibited a heavily infiltrated immune phenotype. Our observations have important implications for prognostication and the development of more effective therapies for this highly lethal variant of bladder cancer.

## INTRODUCTION

Bladder cancer is the ninth most common cancer worldwide, affecting 430,000 people and resulting in 165,000 deaths annually^1^. In the United States, it is the fourth most common cancer in men, with an estimated incidence of 81,000 new cases in 2018^2^. More than 90% of bladder cancers are urothelial carcinomas (UCs), which originate from precursor lesions in the epithelial layer of the bladder, called the urothelium^3^. They progress along two distinct tracks, referred to as papillary and non-papillary, that represent clinically and molecularly different forms of the disease^4^. Non-invasive papillary tumors have a high tendency for recurrence, which necessitates lifetime surveillance that is both intrusive and costly to the patient. Non-papillary carcinomas are clinically aggressive, exhibiting a high propensity for invasive growth, and a large proportion of them are lethal owing to metastatic spread^5^. We showed that papillary tumors are almost exclusively of a luminal molecular subtype that recapitulates the expression pattern of markers characteristic of normal, intermediate, and terminal urothelial differentiation^6^. In contrast, non-papillary UCs are of a basal molecular subtype and exhibit an expression pattern of genes characteristic of the normal basal urothelial layer. Molecular subtyping has shown that invasive UCs can be almost equally divided into luminal and basal subtypes that have distinct clinical behaviors and responses to frontline chemotherapy^6–9^. In addition to conventional UCs, many microscopically distinct bladder cancer variants have been described, and in general they are thought to develop via progression of conventional disease^3,10^. The most frequent of these variants are the sarcomatoid, small cell, and micropapillary, all of which are clinically more aggressive than conventional UCs and require uniquely tailored therapeutic management, which is often unavailable^3,5,10^.

In this report, we focus on one of these more common variants, referred to as sarcomatoid carcinoma (SARC)^3,10^. SARC represents, in various published series, 5-15% of bladder cancer and frequently coexists with conventional UC^11,12^. Clinically, it has a predilection for early metastatic spread to distant organs, and it is associated with shorter survival when compared to conventional UC^10–12^. Here, we report on the genome-wide characterization of bladder SARC, including its miRNA, gene expression, and whole-exome mutational profiles, which identified unique molecular features associated with its aggressive nature that may be relevant for the early detection and treatment of this highly lethal variant of bladder cancer.

## RESULTS

Twenty-eight paraffin-embedded SARC tissue samples and 84 invasive conventional bladder UC samples from the MD Anderson Cancer Center cohort were analyzed retrospectively. A cohort of 408 muscle-invasive bladder cancers in The Cancer Genome Atlas (TCGA) was used as a reference. The samples were characterized by clinical and pathological data as well as by several genomic platforms. Sufficient high-quality DNA was available for 13 SARC cases and 5 paired SARC/conventional UC cases for whole-exome sequencing. Gene expression profiling was performed on all of the cases using Illumina’s DASL platform and the data were merged with those obtained from a cohort of 84 conventional UCs. Panel quantitative reverse-transcription polymerase chain reaction was used to analyze the miRNA expression levels of all 28 SARC samples and 58 conventional UC samples.

### Mutational Signature

The mutational profile of conventional UC was characterized by significant levels of recurrent somatic mutations in 30 genes (**Figure 1A**). The 10 most frequently mutated genes in UC were *TP53* (47%), *ARID1A* (25%), *KDM6A* (22%), *PIK3CA* (22%), *RB1* (17%), *EP300* (15%), *FGFR3* (14%), *STAG2* (14%), *ELF3* (12%), and *CREBBP* (11%). The overall mutational landscapes of luminal and basal bladder UC were similar, but several mutated genes were distinctively enriched in specific molecular subtypes. Mutated *FGFR3*, *ELF3*, *CDKN1A*, and *TSC1* genes were enriched in luminal tumors, whereas mutated *TP53, RB1,* and *PIK3CA* genes were enriched in basal tumors (**Figure 1A**). SARCs showed high overall mutational rates (median mutational frequency 259 with 174 interquartile range) and their significantly mutated genes were similar to those observed in conventional UC (**Figure 1B; Table S1**). However, the top three genes – *TP53* (72%), *PIK3CA* (39%), and *RB1* (39%) – were mutated at significantly higher frequencies in SARCs than they were in conventional UCs (p<0.01). This suggests that SARC evolved from precursor conventional UC carrying these mutations and that mutations in these genes may drive the progression process. Several of the genes that are frequently mutated in conventional UC, including *ARID1A, KDM6A, EP300, ELF3,* and *CREBBP,* were not mutated in SARC, and these genes are involved in chromatin remodeling^13–18^. In general, as a group, chromatin-remodeling genes were not mutated in SARC. Instead, SARC carried frequent mutations of *MYO1F* (33%), *CNGB1* (22%), *FXR1* (22%), *RBM5* (22%), *SEMA3D* (22%), *TTBK2* (22%), and *ZNF90* (22%), which are involved in cellular motility, RNA binding, transmembrane ion channeling, kinase activity, and signaling^19–25^. The functional significance of the mutations of these genes for sarcomatoid progression remains unclear but they are attractive candidates for future mechanistic studies. Interestingly, *FGFR3* mutations, which were present in 14% of conventional UCs, were not present in SARC. Nearly all mutations present in the SARCs were also present in the paired precursor conventional UCs of the same patients, indicating that the SARC and the presumed precursor lesions were clonally related.

**Figure 1.**
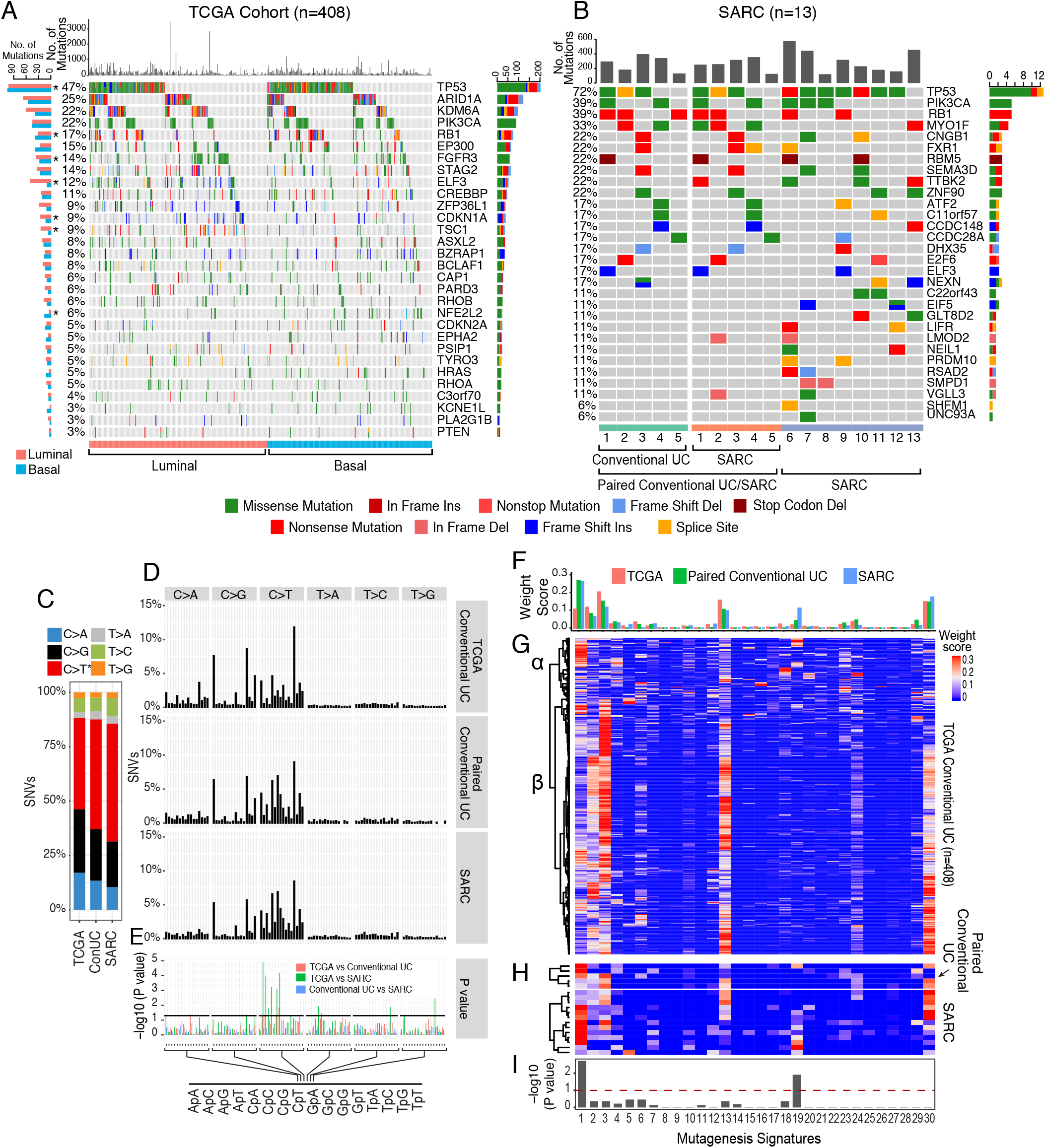
Mutational Landscape of SARC. **(A)** Mutational landscape among the molecular subtypes of 408 muscle-invasive bladder cancers from the TCGA cohort showing the frequency of mutations in individual tumors and somatic mutations for significantly mutated genes. The frequencies of mutations of individual genes in the luminal and basal subtypes are shown on the left. Asterisks denote the genes with statistically significant mutation frequencies in the luminal and basal subtypes. Bars on the right show the numbers of specific substitutions for individual genes. **(B)** Mutational landscape of 13 cases of SARC and 5 paired samples of precursor conventional UC showing the frequency of mutations in individual genes and somatic mutations for significantly mutated genes. The frequencies of mutations of individual genes are shown on the left. Bars on the right show the numbers of specific substitutions for individual genes. **(C)** Composite bar graphs showing the distribution of all nucleotide substitutions in three sets of samples corresponding to the TCGA cohort, paired precursor conventional UC, and SARC. **(D)** Proportion of single-nucleotide variants (SNVs) in specific nucleotide motifs for each category of substitution in three sets of samples as shown in C. **(E)** False discovery rate (FDR) for specific nucleotide motifs in three sets of samples as shown in C. **(F)** Average weight scores of mutagenesis patterns in three sets of samples as shown in C. **(G)** Weight scores of mutagenesis patterns in individual tumor samples of the TCGA cohort. **(H)** Weight scores of mutagenesis patterns in SARCs and paired precursor conventional UCs. **(I)** Statistical significance of mutagenesis patterns in SARC compared with conventional UC.

### Mechanisms of Mutagenesis

To further characterize the mutational process associated with progression from conventional UC to SARC, we examined six single-base substitutions (C>A, C>G, C>T, T>A, T>C, and T>G) in all cancer samples^26,27^. The results revealed that SARCs were enriched with C>T mutations as compared with conventional UCs, and this increase was already apparent in the precursor conventional UCs that were associated with SARCs (**Figure 1C, D**). Analyses of Sanger mutational signatures^28^ revealed the presence of six dominant signatures in the conventional UCs in the TCGA cohort: signatures 1, 2, 3 (BRCA1/2 mutagenesis), 13 (APOBEC), 19, and 30 (**Figures 1F and 1G**). Clustering separated the conventional tumors into two subsets (α and β) that were characterized by different levels of signature 13 (APOBEC) prevalence. In contrast, SARCs and paired precursor conventional UCs were characterized by the uniform dominance of signature 1, which was present in all SARC and precursor conventional UC samples (**Figure 1H**). In addition, clustering segregated SARCs into two subsets, and they were also characterized by different levels of APOBEC activity. Mutagenesis signatures 1 and 19 were significantly enriched in SARCs as compared with conventional UC (**Figure 1I**). Overall, these data reinforce the idea that SARCs evolve from a distinct subset of conventional UCs.

### Gene Expression Profile

Messenger RNA (mRNA) expression profiling revealed that more than 6,000 genes were differentially expressed between SARC and UC. We performed multiple unsupervised clustering analyses using all the differentially expressed genes in SARC and performed similar analyses using the top 100 and top 10 upregulated and downregulated genes. All these analyses separated SARC and UC into two distinct clusters (**Figures S1A and S1B**). One cluster contained conventional UC almost exclusively, whereas the other cluster contained most of the SARCs. Among the top upregulated genes in the SARC cluster were *FAM101B (or RFLNB), UHRF1,* and *PHC2,* all of which are members of the chromatin-remodeling superfamily^29–31^ (**Figure S1B**). The top downregulated genes included the differentiation-associated transcription factor, ELF3, and genes involved in terminal urothelial differentiation, such as uroplakins and cell adherence genes, as well as LAD1, a component of anchoring filaments in the basement membrane^32^. The median survival for the SARC patients (11 months) was significantly shorter than that of the patients with conventional UC (24 months; p=0.0326) (**Figure S1C**).

### Intrinsic Molecular Subtypes

Several molecular studies, including our own, have divided bladder cancer into two molecular subtypes that preferentially express basal or luminal genes^7–9,33^. To investigate whether the intrinsic molecular types of conventional bladder UC applied to SARC, we analyzed the luminal and basal gene expression signatures in SARC. We used a previously developed classifier that includes markers of luminal and basal types^33^. The set of conventional UC was separated into two major groups (**Figure 2A**). The first group, comprising 53 of the 84 samples (63%), was characterized by high mRNA expression levels of luminal markers such as KRT20, GATA3, uroplakins, ERBB2, ERBB3, PPARG, FOXA1, and XBP1 and was referred to as the luminal subtype. The remaining 31 conventional UC samples (37%) were characterized by high expression levels of basal markers such as CD44, CDH3, KRT5, KRT6, and KRT14 and were referred to as the basal subtype.

**Figure 2.**
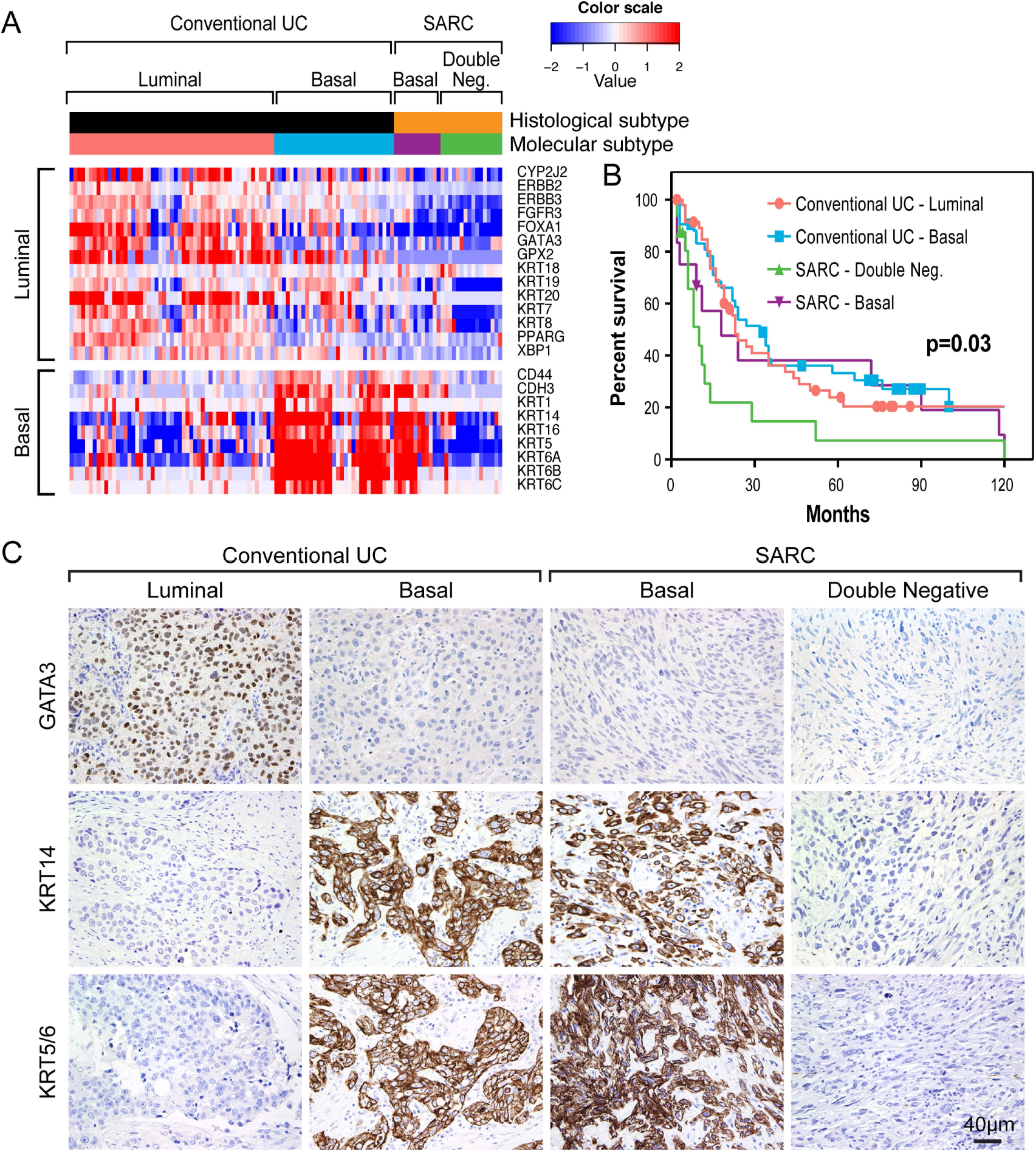
Luminal and Basal Molecular Subtypes in Conventional UC and SARC. **(A)** The expression of luminal and basal markers in molecular subtypes of conventional UC (n=84) and SARC (n=28). **(B)** Kaplan-Meier plots of molecular subtypes of conventional UC and SARC. **(C)** The immunohistochemical expression of signature luminal and basal markers in representative luminal and basal cases of conventional UC and in representative basal and double-negative SARC.

Unlike the conventional UCs, all 28 SARCs had low mRNA expression levels of the luminal genes, which suggested that they developed from a basal subtype of precursor conventional UC (**Figure 2A**). In addition, nearly half of the SARCs (12; 43%) were characterized by the retained expression of basal keratins, CD44, and P-cadherin (CDH3), whereas the remaining SARCs (16; 57%) lacked expression of canonical basal and luminal markers, and we therefore referred to them as “double-negative”. This subset of SARCs shares the “mesenchymal” molecular phenotype with the claudin-low and TCGA cluster IV subtypes identified previously.^7,8,34^ Survival analysis revealed that the double-negative/mesenchymal SARCs were the most aggressive of the molecular subtypes (**Figure 2B**). Furthermore, the mean survival duration of patients with double-negative SARC (10 months) was shorter than that of patients with basal SARC (18 months); however, this difference was not significant statistically, probably because of the limited number of cases. We verified the expression patterns of signature luminal markers (GATA3) and basal markers (P63, KRT5/6, KRT14) by immunohistochemistry using tissue microarrays containing the same cases (**Figure 2C**). The epithelial/basal SARCs were focally positive for basal markers such as p63, KRT5/6, and KRT14 and were negative for the signature luminal marker GATA3. In contrast, the purely mesenchymal double-negative SARCs were immunohistochemically negative for all luminal and basal markers, consistent with the RNA expression data.

### Canonical and Upstream Regulator Pathways

In order to identify candidate mechanisms underlying the progression to the SARC phenotype, we used the Ingenuity Pathway Analysis (IPA) Upstream Pathways and Gene Set Enrichment Analysis (GSEA) functions to identify signaling pathways associated with the gene expression patterns observed. (**Figure 3A-D**) SARCs were characterized by downregulation of G1/S checkpoint genes and upregulation of INK4 family of cyclin-dependent kinase inhibitors (CDKN2A-D), consistent with our observation that *RB1* mutations were more common in SARCs. Similarly, SARCs exhibited decreased p53 pathway activity, which was consistent with their high mutational rate of the p53 gene. SARCs were also characterized by loss of epithelial adherens genes and downregulation of p63 pathway activity complemented with activation of TGFβ and RhoA, all of which are characteristics of EMT permissive state.

**Figure 3.**
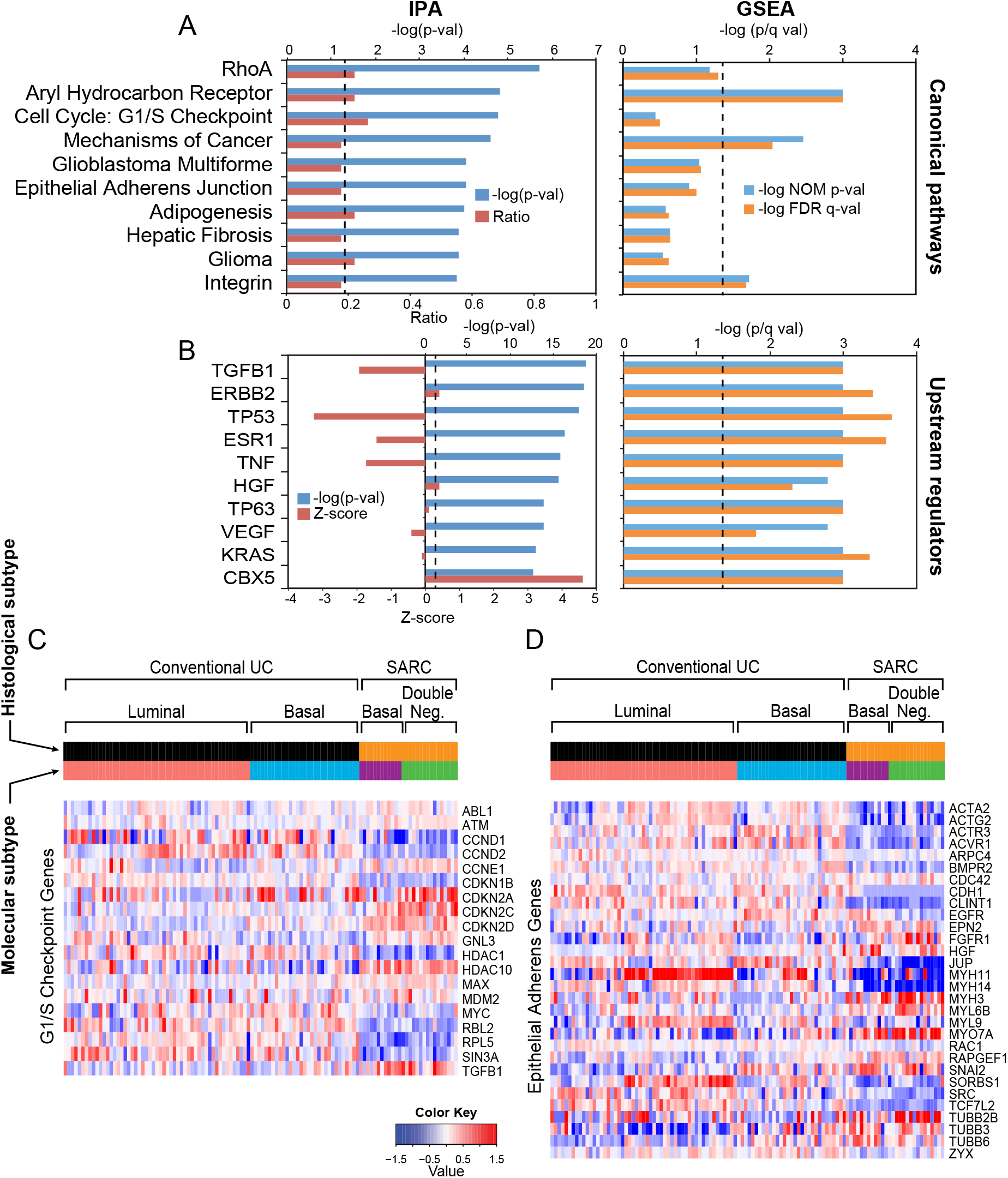
Enrichment Canonical Pathways and Transcriptional Regulators in SARC as Compared with Conventional UC. **(A)** The top 10 canonical pathways dysregulated in SARC, as revealed by IPA and GSEA. **(B)** The top 10 upstream regulators altered in SARC, as revealed by IPA and GSEA. **(C)** Expression patterns of G1/S checkpoint pathway genes in molecular subtypes of conventional UC and SARC. **(D)** Expression pattern of epithelial adherens genes in molecular subtypes of conventional UC and SARC. The dotted lines indicate the significance level.

### Immune Infiltrate

Immune checkpoint blockade is clinically active in about 15% of patients with advanced bladder cancer, where response is associated with high tumor mutational burden (TMB), the “genomically unstable” luminal subtypes, and infiltration with activated cytotoxic T lymphocytes.^35–37^ Given the relatively high mutational frequencies observed in SARCs, we characterized the patterns of immune-related gene expression in them (**Figure 4**). In conventional UCs, 11% (9/84) showed enrichment of an immune gene expression signature and clustered in the luminal molecular subtype. In contrast, 39% (9/28) of the SARCs demonstrated overexpression of immune signature genes and preferentially clustered in the mesenchymal double-negative subtype. The enrichment of immune gene signature in SARC as compared to conventional UC was confirmed by GSEA (**Figure 5A**). We also evaluated the expression signature of immune checkpoint ligands and their receptors^38^ including CD70 and CD27; CD80 and CTLA4; TNFSF9 and TNFRSF18; ADA and ADORA2A; and PDCD1G2 and CD274 (**Figure 5B**). In general, the overexpression of these genes was present in the same group of SARCs characterized by enrichment for the overexpression of immune signature genes. Of potential therapeutic significance, SARC exhibited the enhanced expression of programmed cell death ligand PD-L1 (CD274) and mRNA overexpression of PD-L1 was observed in more than 50% (15/28) of the SARCs (**Figure 5C**). This was confirmed by immunohistochemistry, which showed strong overexpression of the PD-L1 protein (**Figure 5D**), suggesting that immune checkpoint therapy may be an attractive therapeutic option for a subset of SARC patients.

**Figure 4.**
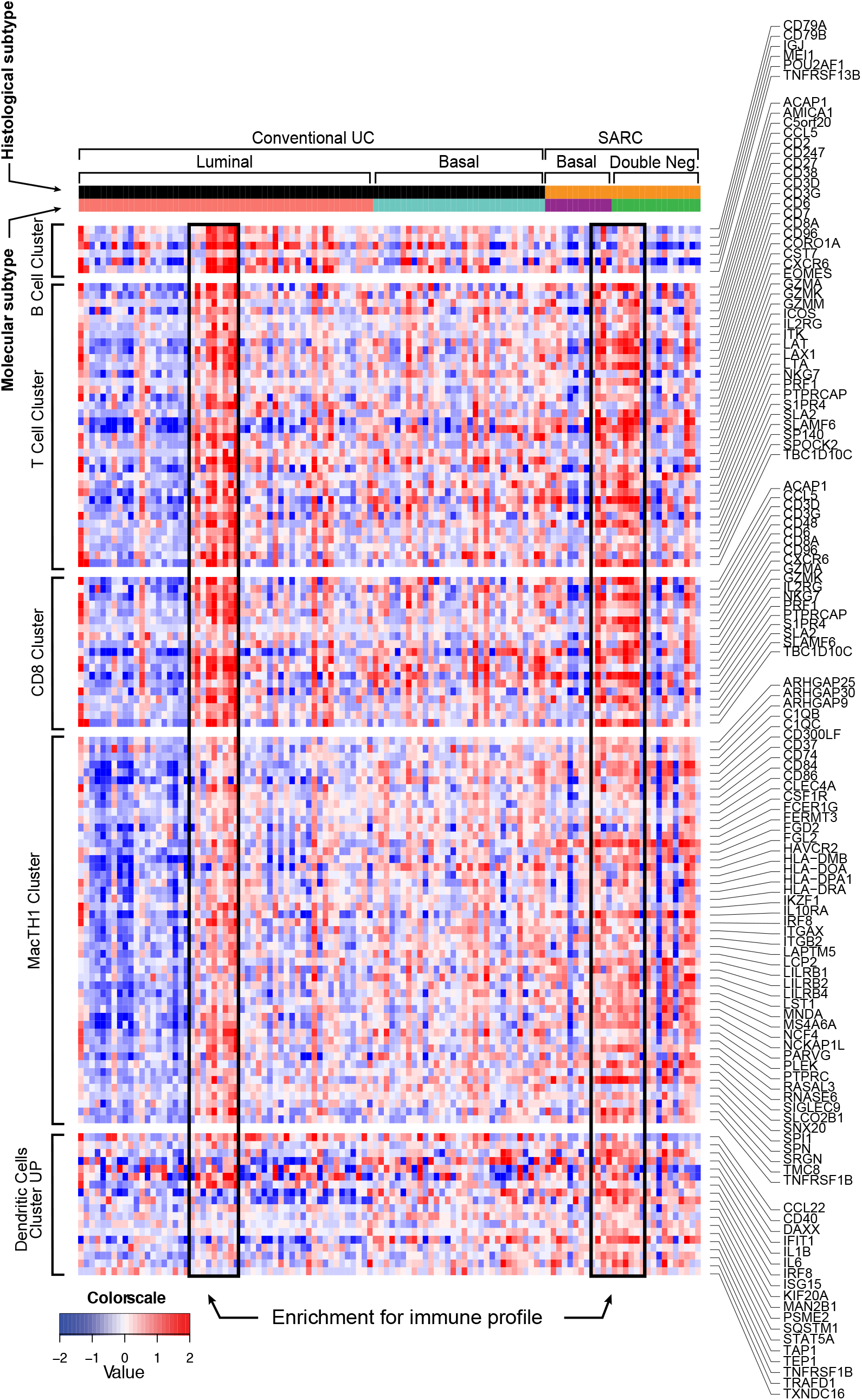
Expression Pattern of Immune Cell Infiltrate in Molecular Subtypes of Conventional UC and SARC. Top to bottom: B-cell, T-cell, CD8, MacTH1, and dendritic cell expression clusters. Boxed areas identify samples with enrichment of immune cell infiltrate.

**Figure 5.**
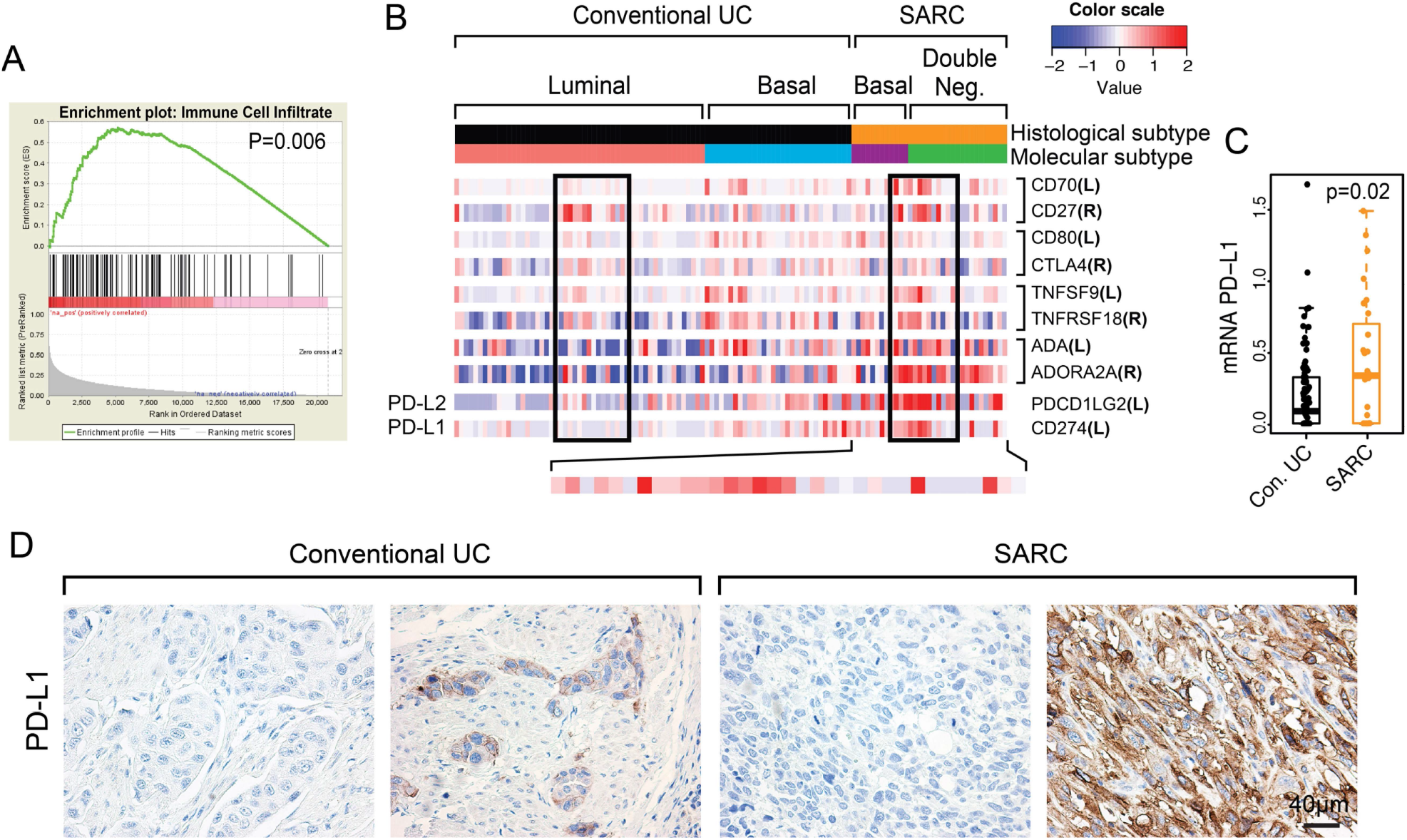
Expression Patterns of Immune Checkpoint Genes in Conventional UC and SARC. **(A)** GSEA of immune cell infiltrate of SARC compared to conventional UC. **(B)** Expression of immune checkpoint genes in conventional UC and SARC in relation to their molecular subtypes. The black boxes indicate the cases with enrichment for immune profile shown in **Figure 4. (C)** Box plot of mRNA PD-L1 expression levels in conventional UC and SARC. **(D)** Examples of immunohistochemical expression of PD-L1 in conventional UC and SARC.

### MicroRNA Expression Profile

Similar to the gene expression profile, microRNA (miRNA) expression levels were widely dysregulated in SARC as compared with conventional UC. Nearly 200 individual miRNA species were either overexpressed or downregulated in SARC (**Figure S2A**). In unsupervised clustering, a small subset of differentially expressed miRNAs could bimodally separate SARC from conventional UC (**Figure S2B**). While the biological significance of the micro RNAs that were upregulated in SARCs is not clear, the entire miR-200 family was downregulated in the double-negative subset of SARCs (**Figure S2B**) and it is likely that it plays an important role in their progression. Members of the miR-200 family play well-established roles in the maintenance of epithelial phenotype by suppressing the transcriptional regulators of EMT of the SNAIL, TWIST and ZEB families.^39^

### Dysregulation of the EMT Network

Microscopically, SARCs comprise mixed lineages of purely mesenchymal cells and cells with at least partial retention of an epithelial phenotype, reflecting various degrees of EMT^40^. We previously showed that TP63 controls the expression of high molecular weight keratins (KRT5, KRT6, KRT14) and suppresses EMT.^6,41^ The central role of p63 in the maintenance of epithelial phenotype and EMT was confirmed in several variants of solid tumors.^42–44^ We therefore performed additional analyses to further characterize the role of EMT in SARC. Among the EMT transcriptional regulators of the SNAIL, TWIST, and ZEB families, SNAIL2 was significantly overexpressed in SARC as compared to conventional UC. Correspondingly, E-cadherin (CDH1) and other epithelial markers including claudin-1 (CLDN1), and tight junction protein 1 (TJP1) were downregulated in SARC (**Figure 6A, Figure S3 and S4A**). In solid tumors, the EMT permissive state may be controlled by p53 and RB1^45^, consistent with the mutational and gene expression characteristics of SARCs. Dysregulation of several additional pathways may have a synergistic effect on the EMT permissive state including the up-regulation of TGFB1 and RhoA, which were among the top up-regulated pathways in SARC by IPA and GSEA. In addition, p63 and its downstream target genes, were downregulated in SARC (**Figures 3A, 3B, and 6A**). Importantly, SARC had widespread downregulation of miRNAs involved in EMT, including miR-100, the p63 targets miR-203 and miR-205, and all members of the miR-200 family^46^ (**Figure 6A and Figure S4B**). Because SARCs can be divided into two subgroups i.e., SARCs that retain epithelial marker expression, reflecting partial EMT, and SARCs that are purely mesenchymal, reflecting complete EMT, we quantified the EMT levels across multiple tumor samples by calculating EMT scores based on a 76-gene signature identified by LA Byers et al.^47^ (**Figure 6B**). This provided a quantitative assessment of the epithelial versus mesenchymal phenotype; positive EMT scores corresponded to the epithelial phenotype, whereas negative EMT scores reflected the mesenchymal phenotype. In general, the conventional UCs were characterized by positive EMT scores corresponding to their epithelial phenotype, whereas the basal and double-negative mesenchymal SARCs had intermediate and low EMT scores, reflecting their partial and complete EMT states. (**Figure 6C**). Essentially identical results were obtained by GSEA using the 175 EMT gene signature developed by M. Yu et al.^48^ The enrichments of SARCs when compared to conventional UCs, as well as SARCs of the basal (partial EMT) subtypes compared to double-negative (complete EMT) were significantly different (**Figures S4C, D**). Immunohistochemistry revealed that tumors within the epithelial SARC subset showed focal retention of p63 and E-cadherin, both of which are involved in the maintenance of epithelial differentiation, whereas double-negative, purely mesenchymal SARCs were negative for the epithelial markers, confirming these subtypes’ partial and complete EMT states, respectively (**Figure 6D**). Finally, the loss of E-cadherin expression was confirmed in selected SARC samples by Western blotting (**Figure 6E**).

**Figure 6.**
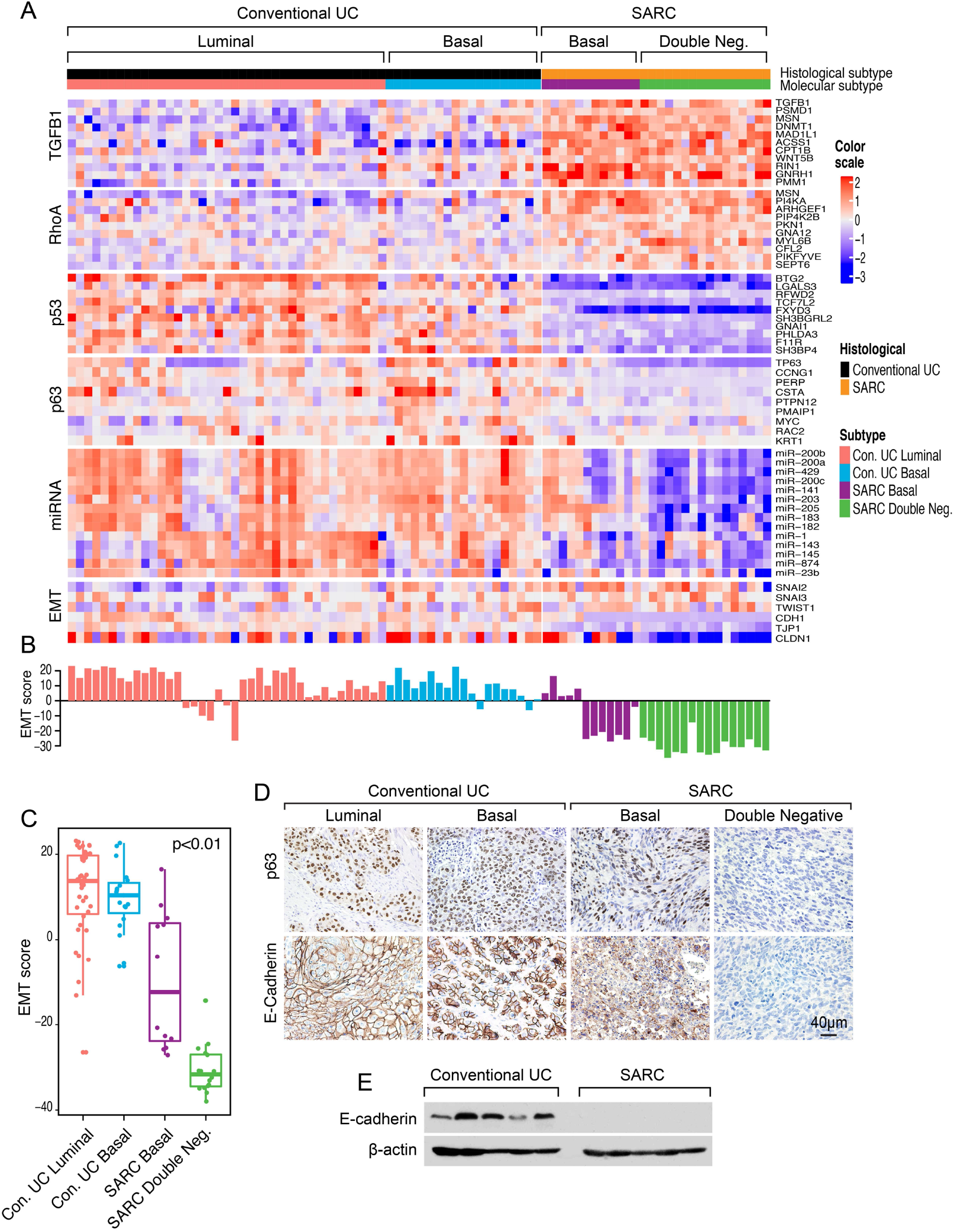
Dysregulation of the EMT Regulatory Network in Molecular Subtypes of Conventional UC and SARC. **(A)** Expression patterns of representative genes in the EMT regulatory network. **(B)** EMT scores in molecular subtypes of conventional UC and SARC. **(C)** Box plot of EMT scores in molecular subtypes of conventional UC and SARC. **(D)** Examples of immunohistochemical expression of p63 and E-cadherin in conventional UC and SARC. **(E)** Western blot documenting loss of E-cadherin expression in SARC.

## DISCUSSION

Bladder cancer is a major source of morbidity and mortality worldwide and in the United States. In the United States, there were approximately 17,000 deaths related to bladder cancer in 2018^2^. Progression to SARC is associated with a high propensity for early metastasis and dismal 5-year survival rates^10–12^. In the following, we highlight essential findings concerning molecular characterizations of SARC and suggest how these findings may contribute to our understanding of its aggressive behavior as well as how they open new therapeutic possibilities.

SARCs have a high overall mutation rate similar to those of conventional UC, melanoma, and non-small cell lung cancers.^49^ Our findings show that in SARC these high mutation rates are associated with mutation signature 1 and that SARCs can be separated into two subgroups, i.e., one with and one without mutation signatures of an endogenous mutagenic enzyme, APOBEC cytidine deaminase. In addition, SARCs are enriched for *TP53, RB1,* and *PIK3CA* mutations as compared to conventional UCs and exhibit gene expression profiles that are consistent with combined loss of TP53 and RB1 pathway activities. In contrast to conventional UCs, and consistent with their basal origins, none of the SARCs profiled here contained activating *FGFR3* mutations, which are enriched in luminal UCs. The absence of mutations in chromatin-remodeling genes is surprising, as SARCs in other organs have been reported to have frequent mutations of these genes, including *ARID1A,* the frequently mutated chromatin-modifying gene in conventional UC^50,51^. On the other hand, the expression analyses have shown that SARCs exhibit upregulation of several chromatin-remodeling genes, including *RFLNB, UHRF1,* and *PHC2,* suggesting that chromatin remodeling gene activity might function to promote EMT, possibly as a cytoprotective mechanism that compensates for their high mutational load^29–31^.

SARCs exhibit a widespread change of their expression profile affecting approximately 30% of the protein-coding genome. However, much of this activity converged on pathways that control EMT, a process in which epithelial cells lose their adhesive features and develop migratory infiltrative properties of the mesenchymal cells^52–54^. The main features of EMT include suppression of E-cadherin and other homotypic adherence and cell polarity genes that is mediated by a group of transcriptional repressors (the SNAIL, TWIST, and ZEB families) complemented by a complex multi-layered regulatory network.^54^ In physiology, EMT is responsible for multi-organ development during the embryogenesis and maintenance or regeneration of these organs in adulthood.^53^ In cancer, including those originating in the bladder, EMT may be a major contributor to the aggressive behavior responsible for invasive growth and metastasis^55^. Our quantitative assessment of EMT showed that basal and double-negative SARCs had intermediate and low EMT scores, respectively, which reflected their partial and complete EMT states, and the purely mesenchymal SARCs were the most aggressive variant of the disease. At the core of this circuitry are p53 and RB, which negatively regulate EMT in solid tumors and appear to be coordinately downregulated in a large percentage of SARCs. Activation of TGFB1 and RhoA can be viewed as a synergistic mechanism complementing the loss of p53 and RB. Another central mechanism that appears to be associated with the double negative SARCs involves downregulation of p63, which positively regulates basal biomarkers (i.e. CDH3, CD44, KRT5, KRT6, and KRT14) expression and negatively regulates EMT via miR-205.^6,41^ In fact, p63 and its target genes expression levels many of which represent basal biomarkers segregate SARCs into epithelial/basal and double-negative purely mesenchymal subtypes. Downregulation of the miR-200 family and other EMT regulatory miRNA species probably reinforces the mesenchymal phenotype of these tumors. Importantly, combined TP53 and RB1 pathways inactivation and up-regulation of EMT appear to be characteristics of the clinically aggressive bladder cancer subset with neuroendocrine phenotype recently identified in the TCGA cohort.^9^ Precisely how these EMT processes are initiated and what are the molecular mechanisms that distinguish sarcomatoid from neuroendocrine progression requires further investigation.

We also found that nearly half of SARCs exhibit an expression signature associated with immune cell infiltration and overexpression of immune checkpoint receptors and their ligands, including PD-L1. This finding suggests new opportunities for immune checkpoint therapy in patients with immune-infiltrated SARCs, which are typically resistant to conventional cisplatin-based chemotherapies^5,11,12^.

On the basis of our findings, we conclude that SARCs are driven by profound dysregulation of the EMT network and that a large proportion of SARCs have an immune infiltration phenotype with upregulation of PD-L1. These features present new avenues of therapeutic potential in patients with this highly lethal variant of bladder cancer.

## METHODS

### Clinical information and tissue samples

All studies and sample collections were performed under Institutional Review Board–approved protocols at MD Anderson Cancer Center. We identified 5,639 cases of bladder cancer, 147 of which were SARC, in a 5-year window from 2008-2013. Most of the SARC cases were outside consultations for which paraffin blocks of tumor tissue were not available. For 28 SARC cases, formalin-fixed, paraffin-embedded (FFPE) tissue was available and sufficient for additional studies. Paraffin blocks from 84 stage- and grade-matched cases of conventional UC were assembled for comparison and clinical data, including patient demographic characteristics, treatments, and outcomes, were retrieved from the patients’ medical records. UCs were classified according to the histologic tumor grading system of the World Health Organization^3^. Microscopically SARCs represented high-grade spindle cell sarcoma in 16 cases and undifferentiated pleomorphic sarcoma in 12 cases. Levels of invasion were defined according to the TNM staging system^56^. All conventional UCs and SARCs were invasive T_2_ and above high-grade tumors. The SARC and UC cohorts had similar age and gender distributions and a male predominance. The mean age of the SARC cohort (22 men and 6 women) was 71 years (range, 41-86 years). The mean age of the conventional UC cohort (65 men and 19 women) was 69 years (range, 33-91 years). The median follow-up times for the SARC and UC cohorts were 9.5 and 23 months, respectively. Together, the two cohorts had at least 90 patients whose deaths were cancer-related. For DNA/RNA extraction and tissue microarray construction histologic slides were reviewed to identify well-preserved tumor-rich areas with minimal amounts of stroma, which were marked on the corresponding paraffin blocks. Four parallel tissue samples were taken from these areas using a 1.0-mm biopsy punch (Miltex, York, PA). In those tumors which contained conventional UC precursor lesions and SARC areas, the two components were sampled separately. Two of the tissue cylinders were used for DNA and RNA extractions for genomic profiling. The other two cylinders were used for the construction of a tissue microarray and for immunohistochemical validation analyses of selected proteins.

### DNA and RNA extraction

Genomic DNA and total RNA were extracted from FFPE tissue samples for DNA sequencing and microarray experiments using the MasterPure Complete DNA and RNA Purification Kit (Epicenter Biotechnologies, Madison, WI) according to the manufacturer’s instructions as previously described^57^. In brief, FFPE tissue cylinders were minced, deparaffinized, and digested with 300_μl Proteinase K digestion buffer with 10_μl Proteinase K (50 μg/μl; Roche Diagnostics, Mannheim, Germany) at 55°C overnight. DNA and RNA concentrations and quality were determined by an ND-1000 spectrophotometer (NanoDrop Technologies Inc., Wilmington, DE) and Quant-iT PicoGreen Kit (Life Technologies, Carlsbad, CA). Sufficient amounts of total RNA for gene expression analysis were extracted from all 28 SARC and 84 conventional UC samples. In addition, sufficient amounts of genomic DNA were extracted from 13 SARC samples, including 5 samples that also contained coexistent precursor conventional UC. DNA extracted from the peripheral blood lymphocytes or normal tissue of the resection specimen from the same patient was used as a control.

### Whole-exome sequencing and processing pipeline

Genomic DNA from 13 cases of SARC and five cases of paired conventional UC were used for whole-exome sequencing, which was performed on the HiSeq 2000 platform (Illumina, San Diego, CA) at MD Anderson Cancer Center’s Genomics Core. The TCGA data on 408 muscle-invasive conventional UCs of the bladder were used as a reference set for mutational analyses. BWA-MEM (version 0.7.12) was used to align reads to the hg19 reference genome. Samtools (version 1.4) and Picard (version 2.5.0) were used to sort and convert between formats and remove duplicate reads^58,59^. The Genome Analysis Toolkit (version 3.4-46) was used to generate realigned and recalibrated BAM files^60,61^. Somatic variants relative to the normal reference sample were detected by MuTect2^62,63^. Oncotator (version 1.8.0.0) was used to produce gene-based and function-based annotations of the single nucleotide variants (SNVs) and insertions/deletions^64^. Similar analyses were performed for the genome-wide expression data from the TCGA cohort (n=408), and tumors were assigned to specific molecular subtypes by applying the sets of luminal, basal, and p53 markers as described previously^33^. Mutational data were downloaded from the TCGA portal (https://tcga-data.nci.nih.gov/tcga/). MutSigCV (version 1.4; https://www.broadinstitute.org/cancer/cga/mutsig) was used to identify genes that were mutated more often than expected by chance given the background mutation processes^65^. The significant gene list was obtained using a false discovery rate (FDR) cutoff of 0.05. The statistical significance of associations between the mutations and the molecular subtypes was assessed by the Fisher exact test.

### Mutagenesis signatures

We used 432 SNVs identified in at least one sample and segregated them into six types of mutations corresponding to the following base pair substitutions: C>A, C>G, C>T, T>A, T>C, and T>G. The Fisher exact test was used to determine the distribution of these mutations in the three groups of samples corresponding to conventional UC in the TCGA cohort and paired UCs and SARCs in the sarcomatoid cohort. The genomic context of SNVs, referred to as fingerprints and which included the two flanking bases on the 5’ and 3’ sides to each position for a total of 96 possible mutational fingerprints, was assembled. Wilcoxon rank sum tests were used to test against the hypothesis of no difference in the frequency of any fingerprint between any two groups of mucosal samples. The Benjamini and Hochberg method was applied to control the FDR. For each sample, we used its mutational fingerprints (V) and the quadratic programming method to compute a weight score *(H)* for each of 30 canonical mutational signatures (W) available from the Sanger Institute (http://cancer.sanger.ac.uk/cosmic/signatures). We applied the 96-by-30 matrix of canonical signatures (W) and, given the 96-by-1 mutational profile of a sample (V), we computed the 30 -y-1 vector (H) for each of the canonical signatures’ relative contributions to the sample profile by solving the following optimization formula:

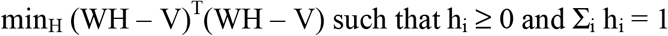

The optimization problems were solved using “quadprog” (version 1.5-5). The Kruskal-Wallis test was used to test against the null hypothesis of no difference in weight scores among conventional UC and paired UC and SARC.

### mRNA expression and data processing

RNAs (0.25–1.0 μg) from SARCs (n=28) and conventional UCs (n=84) were assessed using Illumina HumanHT-12 DASL Expression BeadChips as per the manufacturer’s instructions, and the Illumina BeadStudio v3.1.3 (Gene Expression Module V3.3.8) was used for transformation and normalization of the data. Comparisons were carried out using Welch’s t-tests and Benjamini-Hochberg–controlled FDR–adjusted p-values (<0.05) and fold changes. Unsupervised hierarchical clustering of log ratios was performed with Cluster 3.0, and the results were visualized with Treeview software (Stanford University, Palo Alto, CA). Pearson correlation, mean centering, and average linkage were applied in all clustering applications. Genes within 0.5 standard deviations of the log-transformed ratios were discarded. To select specific and robust gene sets associated with SARC, we used the combination analysis with the Welch t-test and fold-change; genes having p-values <0.05 and showing fold-change >2.0 were selected. IPA software (Ingenuity Systems, Redwood City, CA) was used to determine dysregulated canonical pathways and predicted upstream regulators by calculating z-scores and –log_10_ p-values^66,67^. GSEA was used to evaluate the enrichment probability of the top canonical pathways and upstream regulators identified by IPA^68^. Both SARC and UC samples were classified into luminal, basal, and p53-like intrinsic molecular subtypes using an algorithm described previously^6^.

Immune gene expression signatures for SARC and conventional UC were established using unsupervised hierarchical clustering. Gene dendrogram nodes corresponding to genes characteristically expressed in specific immune cell types were identified and validated through DAVID functional annotation clustering and Ingenuity Systems Analysis (www.ingenuity.com). Immune gene signatures were used as reported previously^69–71^.

To quantitatively assess the level of EMT, we calculated the EMT score based on a 76-gene expression signature reported by Byers et al^47^. For each tumor sample, the score was calculated as a weighted sum of 76 gene expression levels: 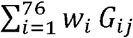, where *w_i_* is the correlation coefficient between the *i*th gene expression in the signature and that of E-cadherin, and is the *i*th gene’s normalized expression in the *j*th tumor sample. We centered the scores by subtracting the mean across all tumor samples so that the grand mean of the score was zero.

### MiRNA analysis

MiRNA analysis was performed on 28 SARC samples and 58 conventional UC samples. For miRNA cDNA synthesis, 400 ng of total RNA was reverse-transcribed using a miRNA reverse transcription kit (Applied Biosystems; catalogue no. 4366596) in combination with the stem-loop Megaplex primer pool (Applied Biosystems). For each cDNA sample, 381 small RNAs were profiled using TaqMan Human MicroRNA A Cards (Applied Biosystems; catalogue no. 4398965). Fold-change for each microRNA was determined using the ΔCt method and examined using Welch’s t-test. An adjusted p-value with FDR < 0.05 was considered significant.

### Validation studies

The expression levels of selected genes were validated on parallel tissue microarrays comprising FFPE samples of 84 UCs and 28 SARCs. The microarrays were designed and prepared as described previously and profiled by genomic platforms^51^. In brief, the tissue microarrays (two 1-mm cores per case) were constructed with a manual tissue arrayer (Beecher Instruments, Silver Spring, MD). Tissue sections from the tissue microarrays were stained with hematoxylin and eosin to confirm the presence of tumor tissue. Immunohistochemical staining was performed with mouse monoclonal antibodies against human GATA3 (HG3-31 clone, 1:100 dilution; Santa Cruz Biotechnology Inc., Santa Cruz, CA), cytokeratin 5/6 (clone D5/16B4, 1:50 dilution; Dako, Carpinteria, CA), cytokeratin 14 (LL002 clone, 1:50 dilution; BioGenex, Fremont, CA), PD-L1 (clone 22C3 *pharmDx* without dilution; Dako), E-cadherin (4A4 clone, 1:1000 dilution; BioCare Medical, Concord, CA), and P63 (4A4 clone, 1:200 dilution; BioCare Medical, Pacheco, CA). Immunostaining was performed using the Bond-Max Autostainer (Leica Biosystems, Buffalo Grove, IL). The staining intensity was scored by two pathologists (CCG and BAC) as negative and mildly, moderately, or strongly positive. In addition, the loss of expression of E-cadherin in SARC was confirmed on selected frozen tumor samples by Western blotting. In brief, whole cell extracts of tumor tissue were analyzed by immunoblotting using an anti-CDH1 antibody (4A, 1:1000 dilution; Cell Signaling Technology, Danvers, MA).

### General statistical analyses

Survival analyses were performed by Kaplan–Meier analysis and log-rank testing. Welch’s t-test was used for two-sample comparison, whereas the Kruskal-Wallis test was used for multiple group comparison. For genome-wide mRNA and miRNA differential expression analysis, the Benjamini and Hochberg (BH) method was applied to control the false discovery rate (FDR). An adjusted p-value with FDR < 0.05 was considered statistically significant.

## Supplementary Figure Legends

**Figure S1. Whole-genome mRNA expression profiling of SARC and conventional UC. (A)** The top 50 upregulated and the top 50 downregulated genes in 28 cases of SARC compared with 84 cases of conventional UC. **(B)** Hierarchical cluster analysis of the cohort shown in **(A)** using the top 10 upregulated and top 10 downregulated genes identified in SARC. **(C)** Kaplan-Meier analysis of survival in SARC and conventional UC.

**Figure S2. miRNA expression profile in SARC and conventional UC. (A)** The top 50 upregulated and downregulated miRNAs in SARC compared with conventional UC. **(B)** Hierarchical cluster analysis of the cohort shown in **(A)** using the top 10 upregulated and downregulated miRNA.

**Figure S3. Dysregulation of the EMT regulatory network in SARC as compared with conventional UC**. Top to bottom, expression patterns of TGFB1, RhoA, p53, and p63 target genes as well as selected miRNA and EMT genes.

**Figure S4. Box plots analysis of genes and miRNAs involved in EMT. (A)** Expression level of genes involved in EMT in TCGA cohort and SARC classified according to the molecular subtypes. **(B)** Expression level of miRNAs involved in EMT in TCGA cohort and SARC classified according to the molecular subtypes. The statistical significance was tested by Kruskal-Wallis rank sum test and is indicated above the individual box plots. All miRNAs tested showed statistically significant difference among the subtypes. **(C)** GSEA for 175 EMT signature genes identified the enrichment for SARC compared to conventional UC in TCGA cohort. **(D)** GSEA for the same group of genes as shown in C displaying the enrichment between basal and double negative/mesenchymal SARC subtypes.

